# PepXPro: a framework for curating, generating, and optimizing structure-affinity protein-peptide datasets

**DOI:** 10.64898/2026.08.09.743757

**Authors:** L. America Chi, F. Marty Ytreberg

## Abstract

Protein-peptide interactions are central to cellular signaling and to a growing class of peptide therapeutics, yet the datasets used to develop and benchmark computational methods for protein-peptide modeling remain poorly standardized. Available databases prioritize comprehensive coverage but require task-specific curation, while published benchmarks are typically distributed as static collections built with heterogeneous curation, quality-filtering, redundancy-reduction, and sampling strategies, limiting reproducibility and cross-study comparison. We present PepXPro, a modular framework that transforms publicly available protein-peptide structure-affinity resources into curated datasets and reproducible benchmark collections generated under user-defined criteria. PepXPro is organized into three components: *Scrape*, for deterministic curation of protein-peptide complex entries from public resources; *GenSample*, for constructing configurable subsets under explicit quality, redundancy, and sampling constraints; and *Benchmark*, for evaluating candidate subsets and selecting a nonredundant, representative, general-purpose benchmark for distribution. Starting from PDBbind and complementary resources, the curation pipeline yields a pool of proteinpeptide complex entries that retains chemically complex cases, including disulfide-linked cyclic peptides, which are commonly excluded from existing benchmarks. We release PepXPro Benchmark v1, a benchmark comprising 70 non-redundant protein-peptide complexes with experimentally determined structures and binding affinities. The underlying framework provides an extensible foundation for reproducible protein-peptide benchmark construction.

## Introduction

Peptide-mediated interactions are fundamental to cellular function, orchestrating diverse biological processes including signal transduction, gene regulation, DNA repair, and immune responses.^1–3^ Moreover, peptides have emerged as attractive therapeutic candidates because of their ability to target large, flat, and shallow interfaces that are often considered “undruggable” by traditional small molecules.^4–6^ This growing interest has driven rapid advances in computational methods for protein-peptide modeling, particularly with the emergence of AI-based methods.^7,8^ The development and fair evaluation of many of these approaches critically depend on high-quality protein-peptide datasets.

Existing protein-peptide resources can be broadly divided into two categories: (i) databases, that prioritize comprehensive data collection and annotation, and (ii) curated subsets that provide datasets optimized for the development, training, or evaluation of computational methods. Existing comprehensive databases differ substantially in scope, ranging from protein-peptide-specific repositories^9–12^ to broader protein-ligand resources,^13^ and vary in the information they provide, including structural data,^9–11^ quantitative binding affinities,^12,13^ and inclusion of search or analysis tools.^12^ However, because their primary objective is comprehensive data coverage, they often contain substantial redundancy, heterogeneous data quality, and uneven sampling across key biological and biophysical properties, requiring additional preprocessing before they can be used for method development or evaluation. The second category comprises curated datasets developed for specific computational applications, including protein-peptide complex structure prediction,^14^ peptide docking, ^15–17^ and structure-based binding affinity prediction.^18^ Many of these resources have become widely used benchmarks for their intended tasks, but they are typically distributed as static datasets generated using fixed quality filters, selection criteria, sampling strategies, and redundancy reduction methods. Consequently, each dataset represents a single realization of the underlying data, reflecting design choices tailored to a specific application while inheriting biases and coverage limitations from the source databases. For example, many datasets exclude chemically complex peptides, such as cyclic peptides, peptides containing non-canonical amino acids, and post-translational modifications, despite their growing importance in peptide-based drug discovery.^5,6^ Likewise, most datasets are biased toward short peptides (typically *<* 15 residues),^15^ with only a few recent resources explicitly incorporating longer and more structurally diverse peptide complexes.^14,16^

Despite these advances, there is currently no widely adopted framework for the automated, reproducible, and extensible generation of standardized protein-peptide datasets that contain 3D structures and binding affinity data. Such a framework is increasingly relevant given the growing use of machine learning methods in protein-peptide modeling and their dependence on high-quality datasets.^7,8,19^ To address this gap, we present PepXPro, an integrated framework for reproducible protein-peptide dataset construction. Specifically, we (i) introduce a deterministic curation workflow for protein-peptide structure-affinity resources; (ii) implement customizable dataset generation under explicit quality, redundancy, and sampling criteria; and (iii) release PepXPro Benchmark v1, a representative, non-redundant dataset that retains chemically complex cases, including disulfide-linked cyclic peptides.

Together, these establish PepXPro as an extensible framework for standardized and reproducible protein-peptide dataset and benchmark generation.

## Methods

PepXPro comprises three complementary components: (1) PepXPro *Scrape*, an automated workflow for downloading, cleaning, filtering, preparing, annotating, and quality-controlling experimentally determined protein-peptide complex structures together with their associated experimentally measured binding affinity data from public databases or user-provided datasets; (2) PepXPro *GenSample*, a flexible framework for generating customized subsets through configurable filtering, sampling and redundancy reduction strategies; and (3) PepXPro *Benchmark*, a procedure used to optimize and generate a non-redundant and representative general purpose benchmark dataset using *Scrape* and *GenSample* modules, culminating in the official PepXPro Benchmark v1 release.

### PepXPro Scrape

#### Input databases

PepXPro uses PDBbind^20,21^ as its primary input source and supports both the original v2020 and refined v2020R1 index releases. Throughout this work, the latest v2020R1 index release is used to illustrate the pipeline. The input consists of entries containing PDB identifiers for experimentally determined protein-peptide complexes together with their associated experimentally measured binding affinities. Additional protein-peptide datasets from external resources (e.g., PEPBI^22^ and PPIKB^12^) or user-provided datasets can also be processed, provided they contain these two required input fields. Regardless of their origin, all entries undergo the same structure preparation, quality control, and metadata annotation workflow to generate standardized datasets. The subset passing this stage is referred to as the *curated* pool.

#### Affinity curation

Binding affinities reported as equilibrium dissociation constants (*K*_d_) or inhibition constants (*K*_i_) are accepted, whereas complexes reported only with IC_50_ values are excluded because these values are not directly comparable to equilibrium binding constants. Binding free energies are computed as follows:

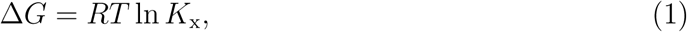

where *R* = 1.987 *×* 10*^−^*^3^ kcal mol*^−^*^1^ K*^−^*^1^ is the gas constant, *T* is the experimental temperature in Kelvin (when reported; otherwise, a default value of 300 K is assumed), and *K*_x_ denotes the reported equilibrium dissociation constant (*K*_d_) or inhibition constant (*K*_i_), expressed in molar units.

#### Structure preparation

Biological assembly mmCIF files are downloaded from the RCSB PDB^23^ for each candidate entry. For structures with multiple biological assemblies, the minimal peptide-receptor interaction unit, defined as the smallest biological assembly containing at least one peptide candidate, is selected. Peptide chains are identified as the shortest protein chains satisfying *L*_pep_ *≤* max(50, 2*×n*), where *L*_pep_ is the length (in residues) of a candidate peptide chain and *n* is the peptide length (e.g., an 8-mer has n=8) obtained from the input metadata when available and otherwise inferred from the peptide sequence. This heuristic tolerates differences between the annotated peptide length and the deposited chain length while excluding large receptor chains from consideration as peptide candidates. Both linear and disulfide-linked cyclic peptides are retained. Receptor chains are defined as protein chains with at least one heavy atom within 4.0 Å of any peptide heavy atom.

Structures were prepared using a custom pipeline built on top of PDBFixer 1.12 and OpenMM 8.5.2.^24^ Preparation includes: (i) removing all water molecules; (ii) retaining only systems with residues and ions that are parameterizable by the AMBER14 force field, with selenomethionine (MSE) converted to methionine (MET); (iii) retaining only AMBER14- compatible ions within 4 Å of the receptor or peptide; and (iv) reconstructing missing residues and side chains, excluding terminal receptor residues (internal gaps only), while reconstructing all missing peptide residues. Entries were additionally rejected if:

- The receptor contains a gap exceeding 10 residues in the deposited structure;
- No receptor chain is found within 4.0 Å of the peptide;
- The peptide is covalently linked to the receptor (terminal C-N distance *<* 1.8 Å);
- The peptide contains fewer than two residues;
- The peptide’s final length (observed residues plus any reconstructed by PDBFixer) exceeds 30 residues;
- Non-AMBER14-compatible residues remain after structure preparation; or
- PDBFixer introduces backbone discontinuities (consecutive C*_α_*-C*_α_* distance *>* 5.0 Å after reconstruction).

#### Metadata

PepXPro generates standardized metadata for every complex containing structural, biophysical, functional, and interface-related annotations. These standardized fields are computed uniformly regardless of the input source, enabling direct comparison across datasets. Metadata native to external resources are preserved and propagated alongside the standardized PepXPro annotations, providing access to dataset-specific information while maintaining a consistent output schema.

Secondary structure for peptides and receptors is assigned using DSSP.^25^ DSSP assignments are reduced to the three-state alphabet (H, E, and C), where H comprises *α*-, 3_10_-, and *π*-helices (H, G, I), E comprises *β*-strands and isolated bridges (E, B), and C comprises turns, bends, and coil conformations (T, S, and unassigned residues). Peptides are then classified as *coil* (no helix or *β*-strand residues present), *mixed* (H *≥* 20% and E *≥* 20%), *helix* (H *≥* 50%), *beta* (E *≥* 50%), *partial helix* (helix present, *<* 50%, helix-dominant), or *partial beta* (strand present, *<* 50%, strand-dominant). This classification was adapted from the secondary-structure categories proposed by Wang et al. ^18^ and Xu et al.^16^

Interface contacts were defined as the number of peptide heavy atoms within 4.5 Å of any receptor heavy atom (*n*_contacts_), whereas hydrogen-bond proxies were defined as interfacial N/O/S atom pairs within 3.5 Å (*n*_hbonds_). In addition, the peptide buried surface fraction is calculated from the solvent-accessible surface area (SASA) using the Shrake-Rupley rolling-sphere algorithm^26^ as implemented in Biopython.^27^ The SASA of the isolated peptide (SASA_free_) and the peptide within the complex (SASA_bound_) are computed independently, and the buried surface fraction is defined as

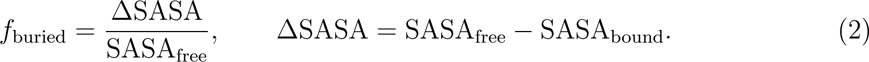

Unlike receptors, peptides generally lack standardized functional annotations. However, their biological context can often be inferred from the functional classification of their cognate receptors. Accordingly, each receptor chain is assigned to one of eight functional classes (immune, enzyme, chaperone, transport, structural, regulatory, signal, peptide-binding, or other) from its PDB entity description. Classification proceeds through a fixed, prioritized set of description keywords, checked in the order above, with the first matching class assigned. Immune is checked first because MHC, HLA, and antibody/Fab chains are not associated with an EC number and would otherwise be misclassified by generic keywords shared with other classes (e.g., “binding protein”, “receptor”). Enzyme classification is further supported by an EC-number check, which captures enzymes identified only by proper name (e.g., trypsin, caspase-7) that no keyword would otherwise match. Entries lacking a usable description, or matching none of the eight keyword sets, are assigned to the “other” category.

#### Implementation details

PepXPro *Scrape* exposes only a small set of user-configurable options, including terminal cap removal and optional preservation of selenomethionine. All other thresholds (e.g., receptor-neighbor cutoff, contact distance, and hydrogen-bond cutoff) are fixed to ensure deterministic and reproducible dataset generation. When integrating external datasets, entries already present in the PDBbind-derived dataset are removed before downloading and processing, ensuring that PDBbind takes precedence. PepXPro *Scrape* produces a standardized summary table containing structural, affinity, and interface metadata, together with processed structures, summary statistics, distribution plots, and optional pairwise receptor TM-score^28^ (template modeling score) matrices for subsequent benchmark generation. TM-score matrices represent length-normalized pairwise global structural similarity between protein structures.

### PepXPro GenSample

PepXPro GenSample applies quality filtering, sequence and structural redundancy reduction, and sampling in a sequential workflow. Each stage is optional and independently configurable.

#### Custom filtering

PepXPro GenSample applies configurable filters to the *curated* pool, including peptide and receptor length, binding affinity range, experimental structure resolution, structure deposition year, peptide topology (linear or disulfide-linked cyclic), and the maximum number of residues modeled to fill missing gaps. The subset passing this stage is referred to as the *eligible* pool.

#### Redundancy reduction

Redundancy is reduced at three complementary levels: receptor sequence, peptide sequence, and receptor structure. Receptor sequences are clustered with CD-HIT,^29^ peptide sequences by pairwise global alignment (Needleman-Wunsch, BLOSUM62, Biopython^27,30,31^), and receptor structures using pairwise TM-score.^32^ These filters can be applied independently or sequentially. The subset passing this stage is referred to as the *non-redundant* pool.

#### Sampling

Following quality filtering and redundancy reduction, the *sampled* subsets are generated using one of several sampling strategies, including stratified, Kennard-Stone,^33^ inverse kernel density estimation (KDE),^34^ and *k*-means clustering.^35^ Sampling can be performed in one or multiple user-defined feature dimensions describing binding affinity, complex size, secondary structure composition, interface properties, and experimental metadata (Table S1 of the Supplemental Material). Different sampling strategies can therefore be selected according to the intended downstream application, such as molecular docking, molecular dynamics, or machine-learning benchmark construction. To support this choice, PepXPro *GenSample* provides a utility that applies all available sampling strategies to the same candidate pool and visualizes the resulting subsets side by side in the selected feature dimensions, allowing users to directly compare how each strategy distributes points across the feature space before committing to a sampling configuration (Figure S1 of the Supplemental Material).

#### Implementation details

PepXPro *GenSample* exposes configurable quality filters, redundancy thresholds, sampling strategies, feature dimensions, and export options. It includes utilities for visualizing dataset statistics and filtering effects, interactively inspecting benchmark structures, and exporting different subsets for downstream applications. Structures may be exported as mmCIF or PDB files, with optional protonation at a user-defined pH. Each run produces a complete parameter record, ensuring full reproducibility.

### PepXPro Benchmark

PepXPro *Benchmark* documents the procedure for generating the official PepXPro Benchmark v1 release. The optimization was performed using the combined PDBbind, PEPBI, and PPIKB datasets to increase the initial number of entries. Candidate entries were restricted to complexes with single-chain receptors to define the *eligible* pool. We next applied a hierarchical three-stage procedure comprising: 1) redundancy configuration selection, 2) sampling configuration selection, and 3) benchmark size selection. Generation parameters were systematically evaluated to identify a default benchmark that combines stringent redundancy filtering with representative, diversity-preserving sampling while maintaining a sufficiently large candidate pool for general-purpose protein-peptide applications. Although this module primarily documents the procedure that was used to produce the PepXPro official benchmark release v1, the framework is fully configurable and can also be applied to generate custom benchmark datasets under user-defined criteria.

#### Benchmark configuration optimization

Redundancy filters were applied sequentially (receptor sequence, peptide sequence, and receptor structure) generating 27 candidate *non-redundant* pools representing all possible redundancy filtering combinations (3 x 3 x 3 = 27 configurations; GC1-GC27)(Table 1). Each candidate pool was first evaluated without sampling using the *Diversity*_pool_ metric, defined as the average of four pool-level metrics describing peptide sequence diversity (PSD), receptor sequence diversity (RSD), receptor structural diversity (RStD), and peptide redundancy fraction (PRF) (Table S3 of the Supplemental Material). Because this metric showed minimal variation across redundancy configurations, the most stringent receptor sequence/peptide sequence/TM-score combination retaining at least 150 candidate complexes was selected, with ties broken in favor of the larger pool (Step 1).

**Table 1:** Parameter space explored during benchmark optimization. Feature-space combinations (D1-D3) and metrics that are used to compute D*_composite_* (M1-M10) and Diversity_pool_ (M5-M7, M10) are detailed in Supplementary Tables S2 and S3, respectively. KS and KDE stand for Kennard-Stone and Kernel Density Estimation, respectively.

| Stage | Parameter | Values |
| --- | --- | --- |
| Redundancy | Receptor sequence (% identity) | strict(40%), medium(50%), lenient(70%) |
|  | Peptide sequence (% identity) | strict(70%), medium(80%), lenient(90%) |
|  | Receptor structure (TM-score) | strict(0.5), medium(0.7), lenient(0.9) |
| Sampling | Method | Stratified, KS, KDE, $k$-means |
|  | Feature space | D1, D2, D3 |
|  | Benchmark size (N) | 25-75% of pool size (5% increments) |
| Diversity evaluation | Diversity score | 4 metrics $\rightarrow Diversity_{\text{pool}}$ |
| Global evaluation | Composite score | 10 metrics $\rightarrow D_{\text{composite}}$ |

Within the selected *non-redundant* pool, different runs were evaluated by varying the sampling strategy, feature space, and benchmark size (Table 1 and Table S2 of the Supplemental Material). Each configuration was scored using the composite metric, *D*_composite_, defined as the average of 10 normalized metrics quantifying feature-space coverage, sequence diversity, structural diversity, redundancy, and distribution preservation, where higher values indicate a better balance among diversity, coverage, and redundancy (Table S3 of the Supplemental Material). All 10 metrics were normalized using bounds computed from runs within the selected redundancy configuration, inverted when lower values indicated greater diversity. The sampling strategy and feature space with the highest mean *D*_composite_ across all benchmark sizes were selected (Step 2). Finally, the benchmark size was defined as the largest sample size whose *D*_composite_ remained within 2% of its maximum value (Step 3).

#### Implementation details

The release was exported with full metadata, fixed mmCIF structures, PDB conversions, peptide and receptor FASTA files, eligible- and redundancy-filtered pool tables, distribution and comparison plots, and a machine-readable parameter record recording the exact command used to generate the release.

## Results

By using the PepXPro *Scrape* module, the initial PDBbind (v2020R1) collection of 19,037 complexes was reduced to 2,458 candidate protein-peptide entries by removing the nonpeptide complexes as a first step. Restricting the dataset to complexes with experimentally determined binding affinities (*K_d_* or *K_i_*) further reduced the dataset to 2,146 entries. An additional 72 complexes were excluded because they contained covalent bonds with the receptor, while subsequent structural quality-control filters-including unsupported AMBER14 residues, the absence of a valid peptide candidate based on length, chain discontinuities, or PDBFixer failures-yielded a final *curated* pool of 708 high-quality protein-peptide complexes with their respective affinities. This *curated* pool comprised complexes spanning a broad range of binding affinities, structural resolutions, deposition years, and complex sizes (Figure 2). Binding free energies ranged from *−*15.5 to *−*2.8 kcal/mol (median *−*7.4 kcal/mol), with most complexes (58%) distributed between *−*9 and *−*6 kcal/mol (Figure 2A). Experimental structures for non-NMR-solved structures (601) had a median resolution of 2.0 Å and 75% of entries resolved at 2.5 Å or better (Figure 2B). The dataset comprises structures deposited between 1994 and 2019, with a median deposition year of 2012 and a marked increase in entries after 2005 (Figure 2C). Peptide lengths ranged from 3 to 19 residues, with a median of 11 residues and 75% of entries containing 14 residues or fewer (Figure 2D).

**Figure 1:**
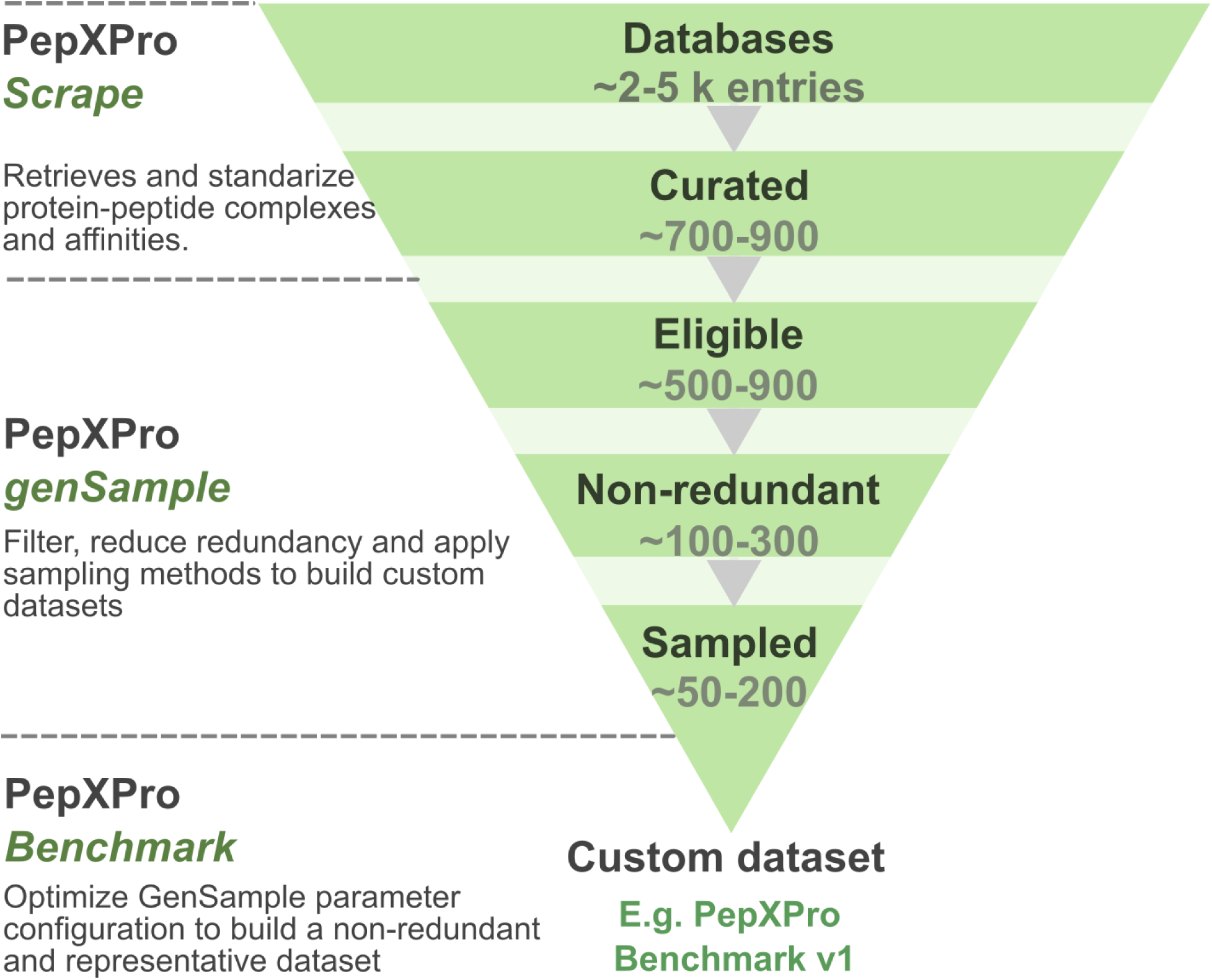
The PepXPro pipeline consists of three modules: *Scrape*, which retrieves and curates protein-peptide complexes from public databases; *GenSample*, which applies user- defined filtering, redundancy reduction, and sampling to generate representative datasets; and *Benchmark*, which optimizes *GenSample* parameters to identify default benchmark configurations. The pipeline enables the generation of custom datasets for different applications, including the PepXPro Benchmark v1 presented in this work.

**Figure 2:**
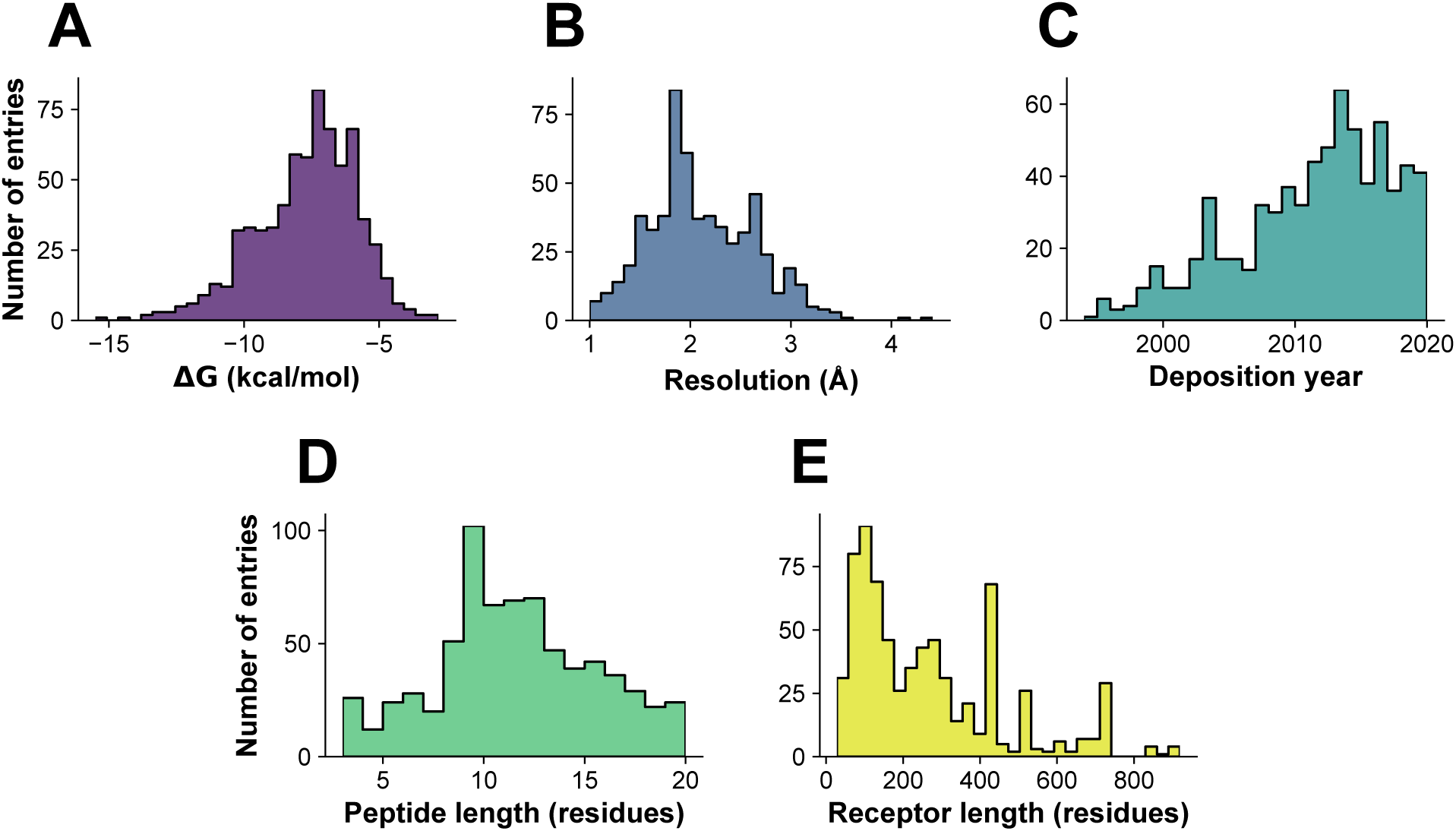
Characteristics of the curated PDBbind dataset using PepXPro (*Curated pool*). Distribution of **(A)** binding free energies (Δ*G*), **(B)** crystallographic resolution, **(C)** deposition year, **(D)** peptide and **(E)** receptor length across the 708 curated protein-peptide complexes.

Receptor lengths showed a broader distribution, ranging from 28 to 919 residues, with a median of 219 residues and 75% of entries containing 375 residues or fewer (Figure 2E). The few receptors shorter than 50 residues (14/708) corresponded to isolated interaction domains deposited experimentally rather than full-length proteins.

Peptide buried SASA fractions ranged from 0.17 to 0.99, with a median of 0.45 (Figure 3A). Interface contacts ranged from 17 to 238, with a median of 56 contacts (Figure 3B). Hydrogen-bond proxies ranged from 0 to 31, with a median of 8 interactions (Figure 3C). Coil peptides constituted the largest secondary-structure class, whereas partial *β*-strand, helical, partial helical, *β*-strand, and mixed conformations were progressively less frequent (Figure 3D). Receptor functional classes were broadly distributed, with enzymes, proteins with no clear category (classified as “other”), signal proteins, and immune-related proteins representing the largest categories (Figure 3E).

**Figure 3:**
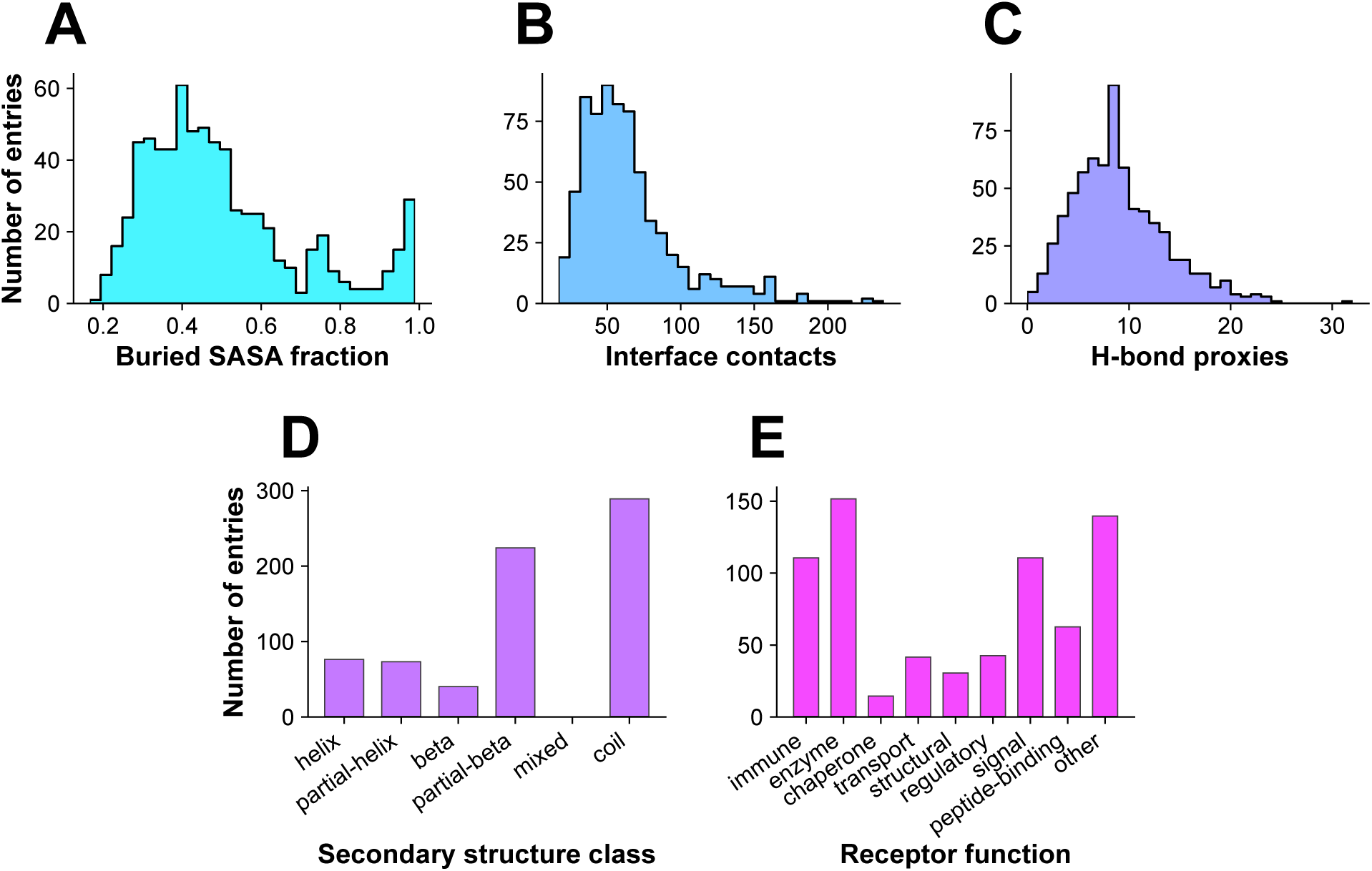
Distributions of **(A**) peptide buried solvent-accessible surface area (SASA) fraction, **(B)** interface contacts, and **(C)** inter-chain hydrogen-bond proxies, **(D)** peptide secondary-structure class, and **(E)** receptor functional class, across the 708 curated protein-peptide PDBbind complexes.

As an illustrative example of the PepXPro *GenSample* module, an arbitrary set of parameter values was used to generate a representative subset of 20 complexes. The curated PDBbind dataset (708 entries) was first restricted to single-chain receptors, producing an eligible pool of 554 complexes. Lenient receptor and peptide sequence redundancy thresholds together with a receptor TM-score cutoff of 0.9 were then applied, reducing the dataset to a non-redundant pool of 244 complexes. Finally, a benchmark of 20 complexes was generated using two-dimensional stratified sampling based on binding affinity (Δ*G*) and peptide length (Figure 4). The resulting benchmark preserved the overall distributions of both variables (Figure 4A,B). Pairwise receptor and peptide sequence identities remained low (*<*50%), with most comparisons below 20%, while receptor structural similarity was also limited, with most pairwise TM-scores between 0.25 and 0.40 and only a few pairs exceeding 0.70 (Figure S2 of the Supplemental Material). The sampled benchmark achieved broad coverage of the binding affinity-peptide length space (Figure 4C), but was not uniformly distributed in the multidimensional feature space (Figure 4D) defined by 12 available descriptors (Table S1 of the Supplemental Material). This example illustrates both the flexibility of GenSample to generate datasets directly under user-defined criteria (no need to pass the optimization stage) and the potential limitations of suboptimal sampling parameters when the goal is broad feature-space coverage, motivating the systematic optimization implemented in PepXPro Benchmark.

**Figure 4:**
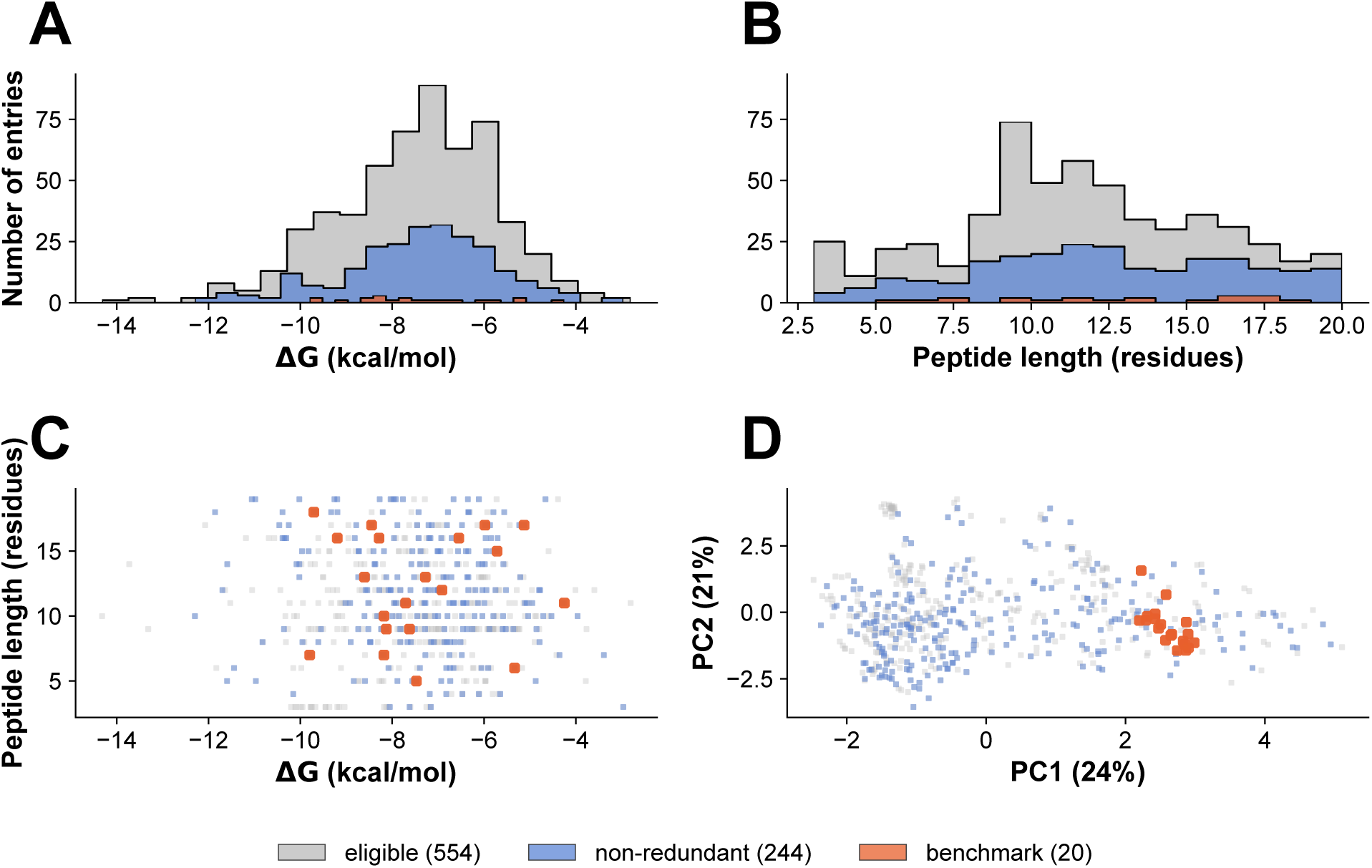
An illustrative benchmark generation run producing a 20-complex benchmark from the PDBbind *curated* dataset. The *eligible* pool (554 complexes) was reduced to a non-redundant pool (244 complexes) using lenient receptor and peptide sequence redundancy thresholds and a receptor TM-score cutoff of 0.9, followed by two-dimensional stratified sampling based on binding affinity (Δ*G*) and peptide length. **(A)** Binding affinity distribution. **(B)** Peptide length distribution. **(C)** Coverage of the Δ*G*-peptide length feature space. **(D)** Principal component analysis of the multidimensional feature space. In all panels, the *eligible* pool, *non-redundant* pool, and *sampled* benchmark are shown to illustrate their relative distribution.

As part of our PepXPro *Benchmark* module, our goal was to generate the largest possible general-purpose benchmark of representative, diverse, and non-redundant protein-peptide complexes with experimentally determined structures and binding affinities. To further expand the coverage of the PDBbind dataset, the same curation workflow was applied to the external protein-peptide resources PPIKB (3,180 original records) and PEPBI (330 original records). Entries whose PDB identifiers were already represented in the curated PDBbind dataset were discarded and, consistent with the PDBbind curation criteria, only structures reporting direct *K_d_* or *K_i_* measurements were retained: 583 and 9 unique candidate structures from PPIKB and PEPBI, respectively. Applying the same structural quality-control pipeline resulted in 187 additional complexes from PPIKB and 5 from PEPBI. Together with the 708 curated PDBbind complexes, these entries produced a merged dataset of 900 protein-peptide complexes (*curated* pool), which served as the source pool for the PepXPro Benchmark v1 release. Despite the incorporation of external resources into the PDBbind dataset, the overall feature distributions remained largely unchanged (Fig. S3-S4 from the Supplemental Material). The principal differences were an extension of the deposition period from 2019 to 2021, an increase in peptide length from 3-19 to 3-30 residues, and the inclusion of larger interaction interfaces with up to 304 residue contacts and a median of 8 hydrogen-bond proxies.

To identify an appropriate redundancy filtering strategy, we evaluated 27 combinations of receptor sequence redundancy (strict, medium, and lenient), peptide sequence redundancy (strict, medium, and lenient), and receptor structural redundancy (TM-score cutoffs of 0.5, 0.7, and 0.9. The merged dataset (900 entries) was restricted to complexes with single-chain receptors, producing an *eligible* pool of 696 complexes. For each configuration, pool-level diversity was quantified using Diversity_pool_ as summarized in the heatmaps of Figure 5A-C alongside each configuration’s resulting pool size. Diversity_pool_ was nearly uniform across the entire parameter space (0.890-0.895), while pool size varied by more than threefold (91-277 entries), indicating that pool-level diversity primarily reflects the severity of the configuration filters. Thus, the redundancy configuration was selected as the most stringent receptor sequence/peptide sequence/TM-score combination whose *non-redundant* pool retained at least 150 complexes, with ties broken toward the larger pool. This procedure selected GC2 (strict receptor redundancy, strict peptide redundancy, TM-score cutoff of 0.7; pool size = 155 entries). For this configuration, Kennard-Stone and cluster-based sampling consistently achieved higher *D*_composite_ values and greater stability across benchmark sizes than KDE or stratified sampling (Figure 5). Kennard-Stone in the D2 feature space (binding affinity, peptide length, receptor length, and interface contacts) achieved the highest mean *D*_composite_ across all tested sample sizes and was selected as the optimal sampling strategy, followed by selecting the largest benchmark size whose *D*_composite_ remained within 2% of the maximum observed value (70 entries).

**Figure 5:**
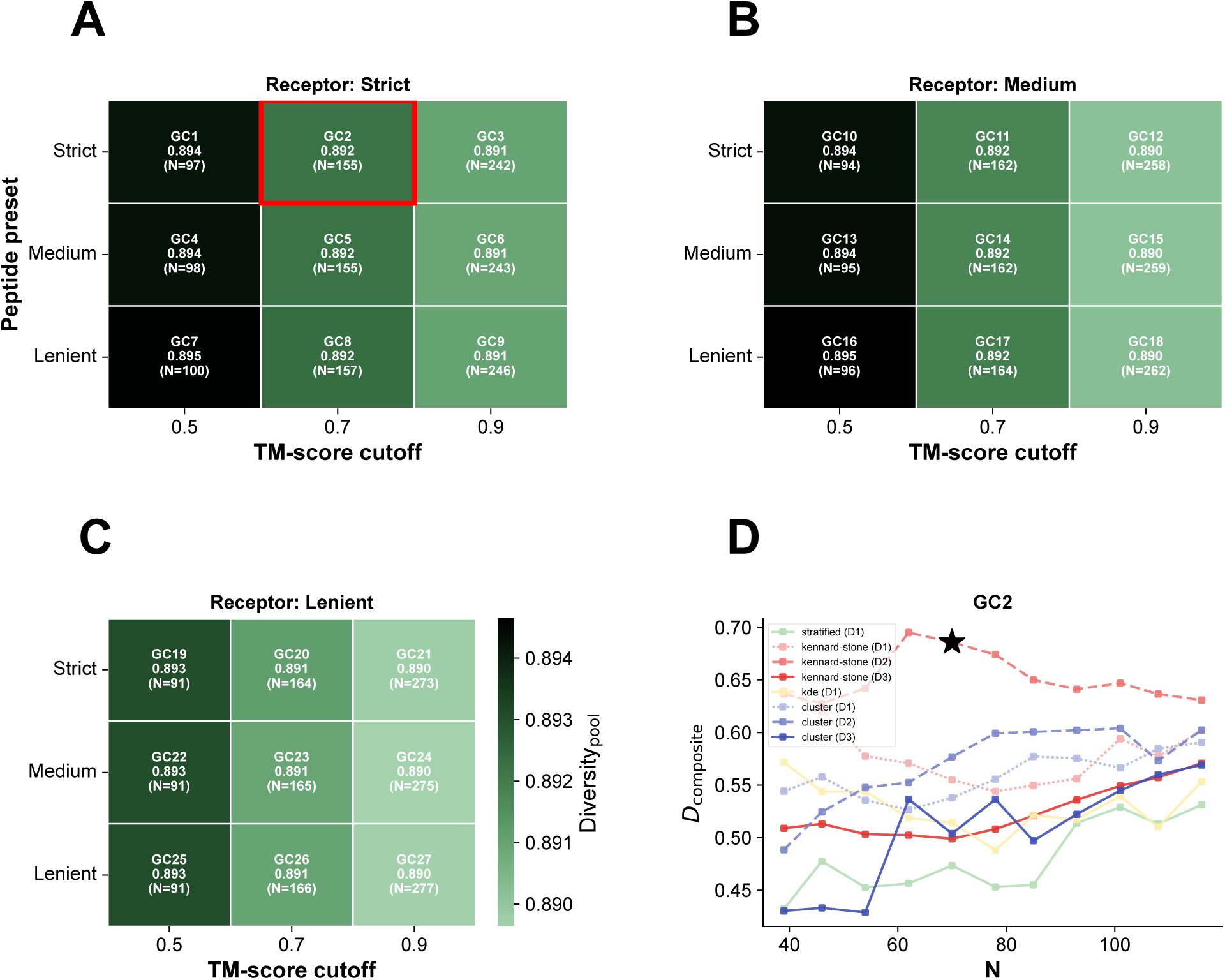
**(A-C)** Pool-level diversity score (Diversity_pool_) for the 27 redundancy filtering configurations generated by combining receptor sequence redundancy (panels), peptide sequence redundancy (rows), and receptor structural redundancy (TM-score cutoff; columns). Each cell reports the grid configuration label (GC1-GC27), its Diversity_pool_ value, and the resulting *non-redundant* pool size (*N*). Darker colors indicate higher Diversity_pool_. The red box highlights the configuration selected as the default for subsequent benchmark generation, chosen as the most stringent filtering combination retaining a sufficiently large candidate pool. **(D)** Detailed performance of the selected configuration (GC2). Unlike panels A-C, which summarize pool-level diversity independent of sampling, panel D shows the composite diversity score (*D*_composite_) as a function of benchmark size (*N*) for each of the four sampling strategies tested (stratified, Kennard-Stone, KDE, and cluster-based). Kennard-Stone and cluster-based sampling were each evaluated in three feature-space combinations (D1-D3), whereas stratified and KDE sampling used only feature space D1. The star in panel D highlights the sample size selected for the final benchmark.

The final benchmark preserved a broad range of structural and biophysical properties while substantially reducing redundancy relative to the initial dataset (Figure 6 and Figures S3-S5 of the Supplemental Material). Sequence redundancy remained low, with median pairwise identities of 12.3% for receptors and 9.1% for peptides, while receptor structural similarity was also limited (median TM-score = 0.32)(Figure S5 of the Supplemental Material). Binding affinities spanned from *−*13.7 to *−*3.0 kcal/mol (median = *−*7.4 kcal/mol)(Figure 6A), peptide lengths ranged from 3 to 30 residues (median = 14)(Figure 6B), and receptor lengths ranged from 33 to 902 residues (median = 191)(Figure 6C). Similarly, complexes exhibited between 17 and 286 receptor-peptide interface contacts (median = 61.0), reflecting a wide spectrum of interaction sizes (Figure 6D). Coverage is representative across the four features sampled in the sampling stage (affinity, peptide length, receptor length, and interface contacts)(Figure 6E-F). Principal component analysis indicated that the benchmark broadly sampled the multidimensional feature space of the original pool, with minimum nearest-neighbor distances ranging from 1.11 to 3.65 standardized units (median = 2.05), indicating good coverage without excessive clustering (Figure 6G). The benchmark also retained diverse values of buried SASA fractions, *n*_hbonds_, peptide secondary structures (Figure 6G and receptor functional classes, suggesting that the selection strategy successfully balanced structural, sequence, and functional diversity (Figure 7).

**Figure 6:**
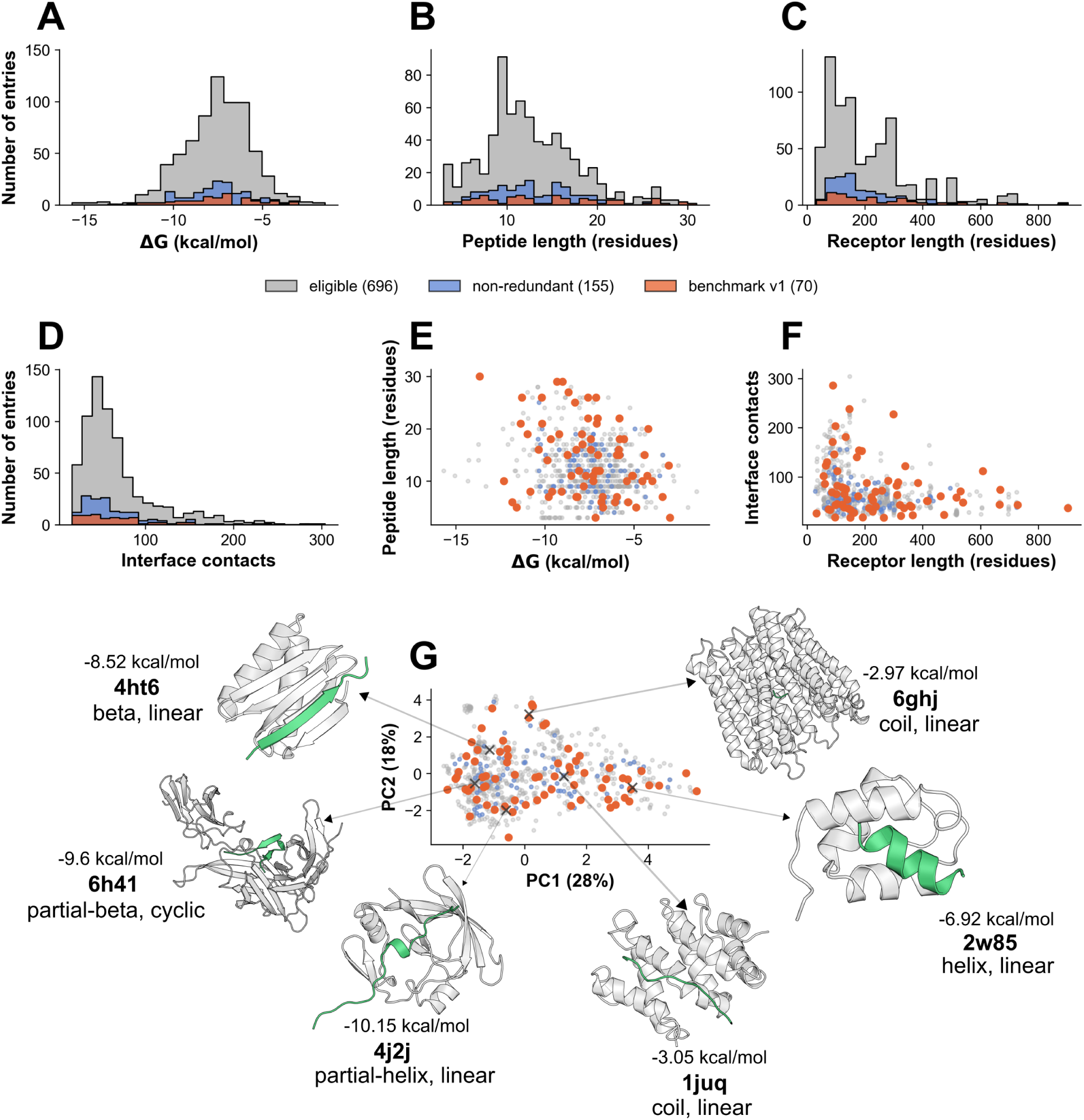
Distribution of **(A)** binding affinities (Δ*G*), **(B)** peptide lengths, **(C)** receptor lengths, and **(D)** interface contacts for the *eligible* dataset (gray), *non-redundant* pool (blue), and final *sampled* benchmark (orange). **(E)** Relationship between binding affinity and peptide length. **(F)** Relationship between receptor length and the number of receptor-peptide interface contacts. **(G)** Principal component analysis of the multidimensional feature space, showing that benchmark complexes are broadly distributed throughout the non-redundant pool. Six representative protein-peptide complexes selected to illustrate the structural diversity of the PepXPro v1 benchmark release. The examples include helical peptides (4j2j, 2w85), *β*-strand peptides (6h41, 4ht6), an extended coil peptide (1juq, 6ghj), and a cyclic peptide (6h41). Receptors are shown as gray cartoons and peptides in green. Experimental binding free energies (Δ*G*) and PDB identifiers, secondary structure class and topology for the peptides are shown below each structure.

**Figure 7:**
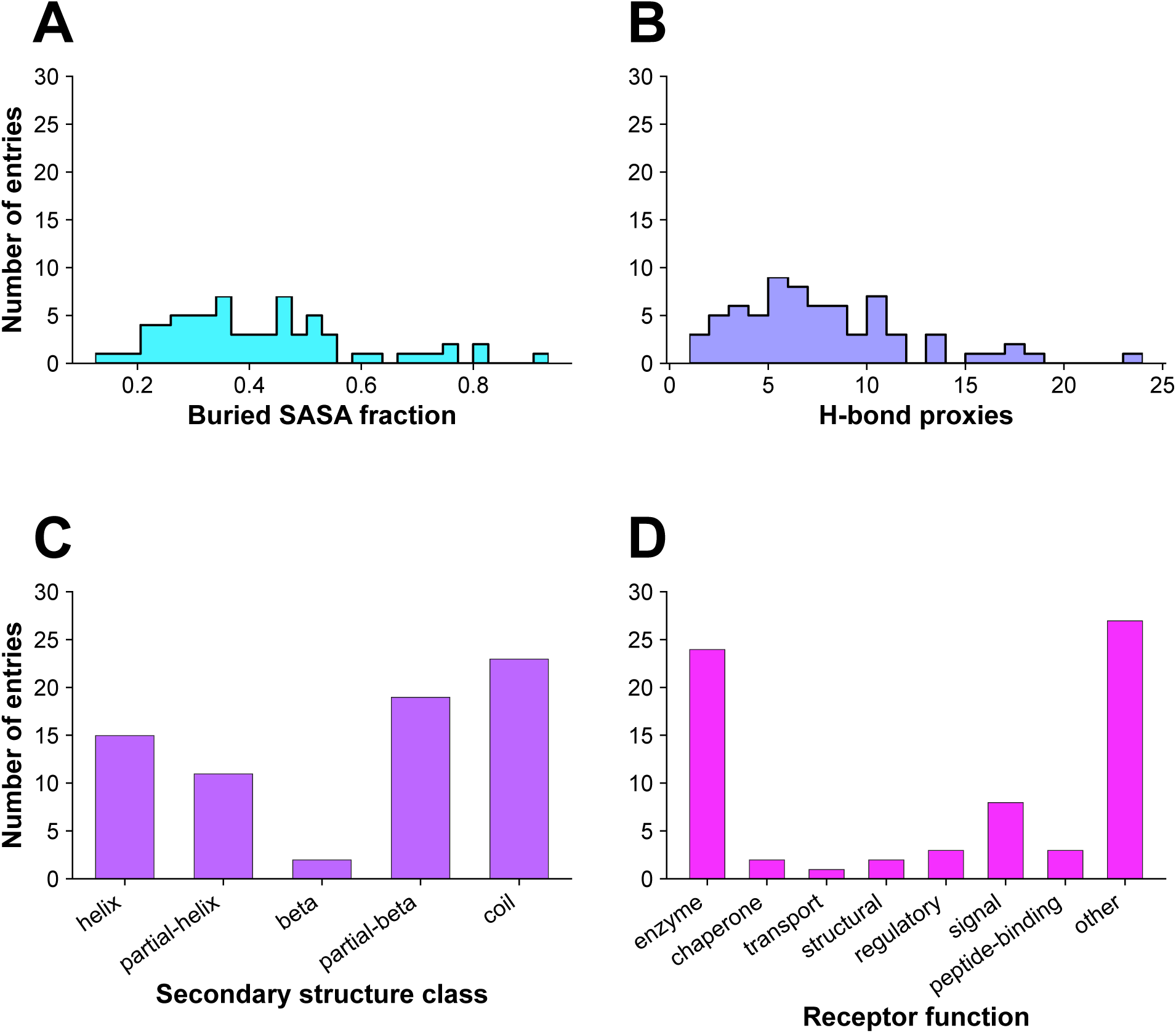
PepXPro Benchmark v1 release. **(A)** Distribution of buried SASA fractions. **(B)** Distribution of *n*_hbonds_. **(C)** Distribution of peptide secondary structure classes. **(D)** Distribution of receptor functional categories in the final benchmark.

## Discussion

PepXPro occupies a distinct position within the landscape of protein-peptide resources. Existing databases such as PepBDB, ^9^ PepBind,^10^ Propedia,^11^ PDBbind^20,21^ and PPIKB^12^ prioritize comprehensive coverage and annotation of entries, but their heterogeneous quality and systems redundancy require substantial preprocessing before they can be used for method development or evaluation. Conversely, curated benchmarks such as LEADS-PEP,^15^ PepPro,^16^ and the structure-based affinity set of Wang et al.^18^ provide task-specific, nonredundant collections. Each of these collections is distributed as a single static realization built with fixed filtering, sampling, and redundancy-reduction choices, and most are biased toward short linear peptides while excluding chemically complex ones. PepXPro differs from both categories in three respects: (i) it curates experimentally determined structures with experimentally measured binding affinities under a single deterministic, reproducible curation workflow; (ii) it generates subsets or benchmarks on demand under user-configurable quality, redundancy, and sampling criteria rather than shipping a fixed collection; and (iii) it retains cyclic (disulfide-linked) and large (15-30 residues) peptides that are commonly discarded by existing benchmarks. Unlike frameworks aimed at evaluating a specific downstream task, such as PepPCBench^14^ designed for protein-peptide complex-structure prediction, PepXPro targets the upstream problem of standardized, extensible dataset construction, providing a common foundation on which a broader range of task-specific benchmarks can be built.

The strengths and limitations of PepXPro are ultimately determined by the underlying experimental datasets from which it is built. PDBbind is not exclusively composed of protein-peptide complexes and contains substantial heterogeneity in its data, making the identification and curation of bona fide protein-peptide entries a major challenge. To address this issue, the PepXPro *Scrape* module provides a standardized, automated, and reproducible curation workflow that minimizes subjective manual intervention. Despite this systematic curation, the resulting dataset reflects the current limitations of available experimental data. The reduction from nearly 20,000 PDBbind entries to only 708 curated protein-peptide complexes highlights the limited availability of experimentally resolved peptide-based complexes with experimentally determined binding affinities suitable for benchmarking. Likewise, the predominance of intermediate binding affinities, structures deposited before 2020, short peptides, enzyme targets, proteins represented by isolated binding domains rather than fulllength structures, and coil-rich peptide conformations reflects biases inherent to the current structural landscape rather than limitations introduced by PepXPro. Nevertheless, the PepXPro benchmark v1 release demonstrates that is possible to generate representative subsets that retain substantial diversity in affinity, structural quality, size, and deposition time, providing a robust foundation for benchmark optimization.

An effective protein-peptide benchmark must satisfy several widely recognized design principles: non-redundancy, to prevent inflated performance from structurally or evolutionarily similar train-test cases; diversity, spanning peptide length, secondary-structure content, and biological function; chemical complexity, so that non-canonical and cyclic peptides are represented rather than discarded; representativeness of the real-world distribution of binding affinities and interface properties; and standardized, tool-independent construction through transparent, reproducible pipelines. PepXPro partially addresses these principles. Nonredundancy is enforced at three levels (receptor sequence, peptide sequence, and receptor structure); diversity preservation is assesed through the composite score together with the distributions of peptide secondary structures and receptor functional classes; chemical complexity is preserved by retaining disulfide-linked cyclic peptides that are commonly excluded from existing sets while more challenging larger peptides are also retained; representativeness is assessed by comparing feature distributions across the eligible, non-redundant, and benchmark levels; and standardization is achieved through a deterministic, fully documented pipeline that is independent of any particular predictive method it may be used to evaluate. Temporal independence is likewise supported at the framework level: GenSample provides a deposition-date filter that restricts candidate complexes to a user-defined release window, enabling the construction of time-split benchmarks that are unseen by models trained up to a given cutoff. In practice, however, this capability is currently constrained by the source data rather than the framework, since PDBbind, the primary resource of PepXPro, does not include post-2019 structures, and the few later entries from complementary resources are insufficient to enforce a meaningful time split. Consequently, the present release cannot yet guarantee temporal independence from contemporary structural databases, a limitation that reflects the state of the available affinity-annotated data and that will be relaxed as newer structure-affinity resources are incorporated.

There is an inherent trade-off between dataset representativeness, feature-space coverage, redundancy reduction, and diversity preservation. Preserving the distributions of experimentally observed properties, such as binding affinity and peptide length, promotes representativeness, whereas preservation of diversity and reduction of redundancy favor the selection of more distinct complexes. Similarly, maximizing feature-space coverage encourages the inclusion of entries from sparsely populated regions of the dataset. Consequently, no single sample-generation strategy is universally optimal; rather, the appropriate balance depends on the intended downstream application. For example, machine-learning applications generally benefit from representative benchmarks that closely match the underlying data distribution, whereas docking benchmarks, blind prediction challenges, or method validation may benefit from maximizing structural and sequence diversity to reduce redundancy and broaden the explored conformational space. Rather than enforcing a single definition of an optimal benchmark, PepXPro exposes these trade-offs through configurable redundancy thresholds, feature spaces, sampling methods, and benchmark sizes, allowing users to tailor benchmark construction to the requirements of specific computational tasks. The official PepXPro Benchmark v1 should be regarded as a well-balanced reference configuration rather than a definitive solution.

Several limitations should be considered when using PepXPro. First, the affinity data are inherently heterogeneous: dissociation (*K*_d_) and inhibition (*K*_i_) constants are pooled into a single Δ*G* scale via Eq. 1, even though *K*_i_ is not strictly thermodynamically equivalent to *K*_d_, and complexes reported only with IC_50_ values are excluded altogether, discarding a substantial fraction of otherwise usable entries. In addition, when the experimental temperature is not reported, a default of 300 K is assumed, introducing a systematic uncertainty in the derived Δ*G*. Second, structural coverage is constrained by the force-field-based preparation pipeline: only residues and ions parameterizable by AMBER14 are retained, so peptides containing non-canonical amino acids or post-translational modifications are removed, and only disulfide-linked cyclic peptides are kept, excluding head-to-tail and other macrocyclic topologies. This is a notable constraint given that these chemically complex peptides are of growing importance in peptide-based drug discovery.^5,6^ Finally, the official PepXPro Benchmark v1 was restricted to single-receptor-chain complexes and therefore does not represent multi-chain or higher-order interaction interfaces. However, users can choose to include such systems in their own benchmarks if desired. The modular, user-friendly, and open-source design of PepXPro, together with its implementation in Python, provides an extensible framework that can progressively address the above-mentioned limitations through the integration of additional data sources and dataset preparation options in future releases.

Continued generation and deposition of diverse, high-quality affinity measurements for complexes with experimentally determined structures will therefore be essential, both to expand benchmark coverage and to support the rigorous evaluation of future predictive methods.

## Conclusion

We introduced PepXPro, a modular framework that converts publicly available protein-peptide structure-affinity resources into curated datasets and reproducible, on-demand benchmark collections through its Scrape, GenSample, and Benchmark modules. Applied to PDB- bind and complementary resources, the pipeline yields a curated pool that retains chemically complex cases, such as disulfide-linked cyclic peptides, that are commonly excluded from existing benchmarks, and it exposes the quality, redundancy, diversity, and sampling choices underlying a benchmark as explicit, user-controlled parameters. Using this framework, we generated PepXPro Benchmark v1, a non-redundant, representative release of 70 protein-peptide complexes with experimentally determined binding affinities. Beyond this specific release, the framework provides a transparent and extensible foundation for constructing standardized protein-peptide benchmarks, enabling reproducible dataset generation and fair cross-study comparison as new structural and affinity data become available.

## Acknowledgement

This work was supported by the U.S. Department of Defense (DOD) under project 1I80VP000634-01. Computational resources were provided in part by Research Computing and Data Services in the Institute for Interdisciplinary Data Science at the University of Idaho.

## Author contributions statement

FMY conceived and conceptualized, and supervised the project. FMY and LAC developed and implemented the workflow. FMY and LAC analyzed the results. The initial draft of the manuscript was prepared by LAC, and all authors contributed to reviewing and editing the manuscript.

## Data Availability

The PepXPro framework is distributed as open-source software at https://github.com/YtrebergLab/PepXPro, together with the curated dataset and the PepXPro Benchmark v1 release and a step-by-step tutorial and a user manual describing the Scrape, GenSample, and Benchmark modules and their configuration options.

