## Supplemental Information for "PepXPro: a framework for curating, generating, and optimizing structure-affinity protein-peptide datasets"

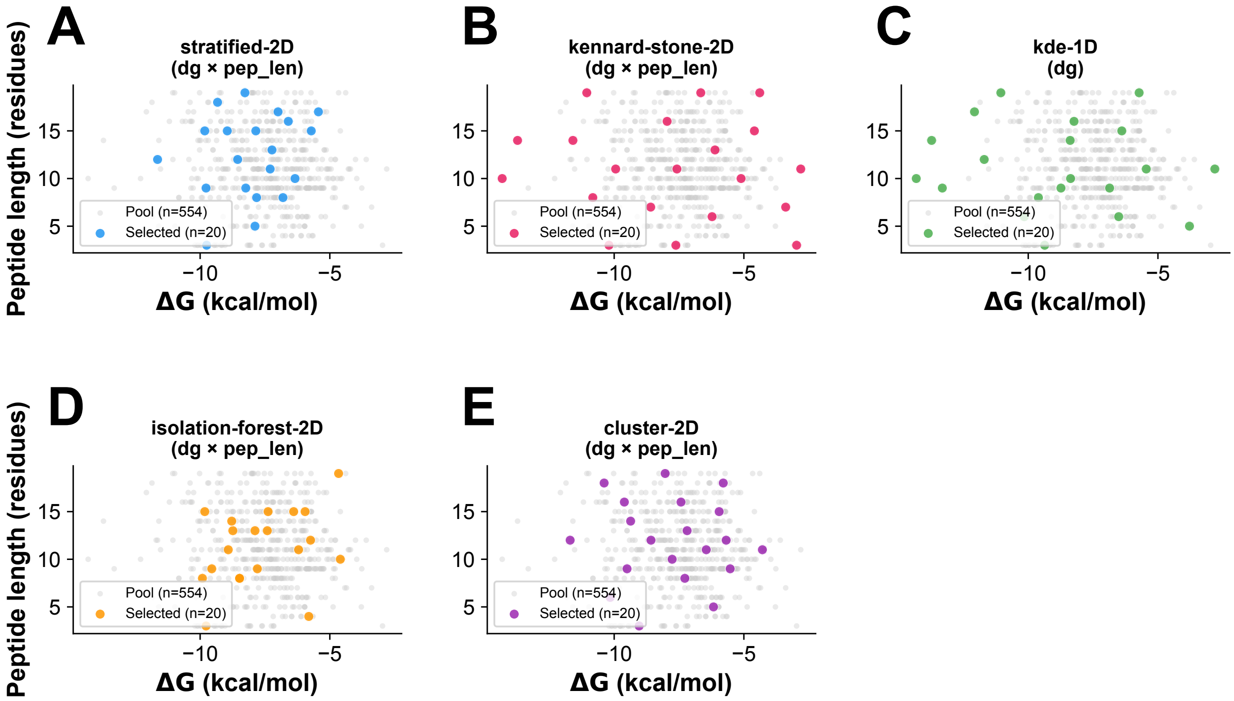


Figure S1: **Comparison of available sampling strategies.** Representative subsets of 20 complexes were generated from the same eligible pool (555 complexes) using 2D stratified sampling, 2D Kennard-Stone sampling, 1D kernel density estimation (KDE), 2D isolation forest, and 2D clustering. The *eligible* pool comes from the *curated pool* from PDBbind only and is filtered to single chains only to generate the *eligible* pool. Gray points represent the *eligible* pool, and colored points denote the selected complexes. Sampling was performed using binding free energy ($\Delta G$) and peptide length, except for KDE, which used only $\Delta G$.

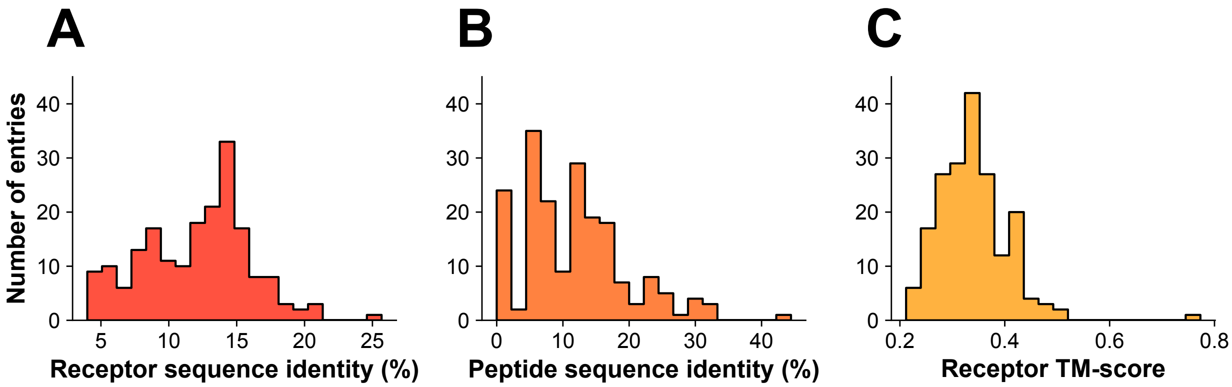


Figure S2: An illustrative benchmark generation run producing a 20-complex *sampled* subset (run_01) from the curated PDBbind dataset. The *eligible* pool (554 complexes) was reduced to a *non-redundant* pool (244 complexes) using lenient receptor and peptide sequence redundancy thresholds together with a receptor TM-score cutoff of 0.9, followed by two-dimensional stratified sampling based on binding affinity ($\Delta G$) and peptide length. (A) Distribution of receptor sequence identity within the benchmark, (B) distribution of peptide sequence identity, and (C) distribution of receptor structural similarity (TM-score).

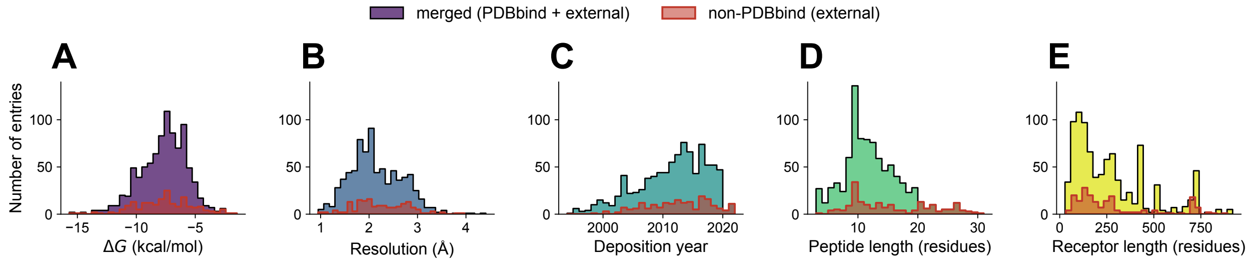


Figure S3: Characteristics of the curated PDBbind + external dataset (PPIKB and PEPBI) by using PepXPro. Distribution of (A) binding free energies ($\Delta G$), (B) crystallographic resolution, (C) deposition year, (D) peptide length, and (E) receptor length across the 900 curated protein-peptide complexes.

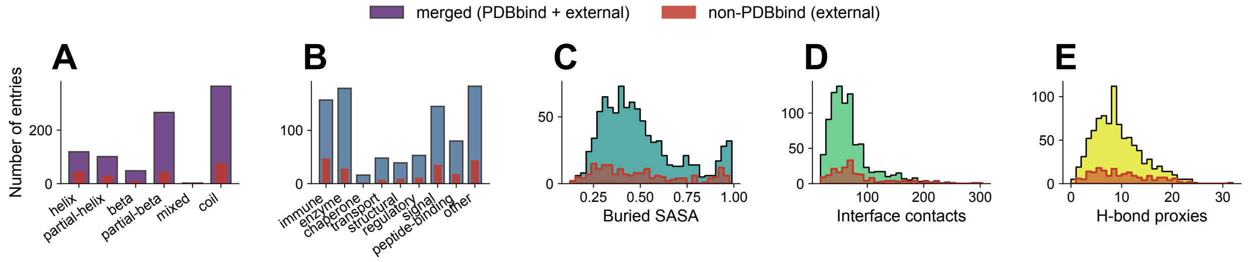


Figure S4: Distributions of (A) peptide secondary-structure class, (B) receptor functional class, (C) peptide buried solvent-accessible surface area (SASA) fraction, (D) interface contacts, and (E) inter-chain hydrogen-bond proxies across the 900 curated protein-peptide complexes from PDBbind + external dataset (PPIKB and PEPBI).

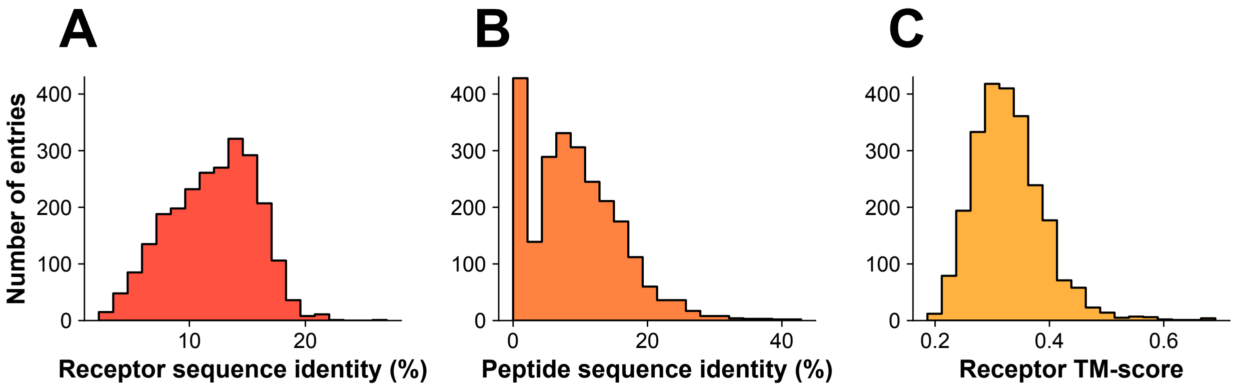


Figure S5: Benchmark v1 release. (A) Distribution of receptor sequence identity within the benchmark, (B) distribution of peptide sequence identity, and (C) distribution of receptor structural similarity (TM-score).

Table S1: Features available for constructing the multidimensional feature space used by PepXPro *GenSample*, encompassing both the fixed set used for principal component analysis (PCA) and the full set of features that may be selected for subsampling.

| Feature | Description | PCA |
|---|---|---|
| `dG` | Binding free energy ($\Delta G$) | ✓ |
| `pep_len` | Peptide length (residues) | ✓ |
| `rec_len` | Receptor length (residues) | ✓ |
| `n_rec_chains` | Number of receptor chains | |
| `ss_class` | Peptide secondary-structure class (categorical: helix, partial-helix, beta, partial-beta, mixed, coil, other) | |
| `func_class` | Receptor functional class (categorical: immune, enzyme, chaperone, transport, structural, regulatory, signal, peptide-binding, other) | |
| `pep_helix` | Peptide $\alpha$-helix content (%) | ✓ |
| `pep_beta` | Peptide $\beta$-strand content (%) | ✓ |
| `pep_coil` | Peptide coil content (%) | ✓ |
| `rec_helix` | Receptor $\alpha$-helix content (%) | ✓ |
| `rec_beta` | Receptor $\beta$-strand content (%) | ✓ |
| `rec_coil` | Receptor coil content (%) | ✓ |
| `n_contacts` | Number of protein–peptide interface contacts | ✓ |
| `n_hbonds` | Number of protein–peptide hydrogen bonds (proxy) | ✓ |
| `sasa_burial` | Buried solvent-accessible surface area (SASA) of the peptide | ✓ |
| `binding_kd` | Binding measurement type indicator ($1 = K_d$, $0 = K_i$) | |
| `resolution` | Crystallographic resolution (Å) | |

Table S2: Feature-space combinations evaluated during benchmark optimization.

| ID | Feature dimensions |
|---|---|
| D1 | Binding affinity ($\Delta G$), peptide length |
| D2 | D1 + receptor length, interface contacts |
| D3 | D2 + peptide secondary structure (helix, $\beta$-sheet, coil), receptor secondary structure (helix, $\beta$-sheet, coil), buried SASA |

Table S3: Metrics used to compute the composite score ($D_{\text{composite}}$) and the whole-pool diversity score ($\text{Diversity}_{\text{pool}}$). Metrics are labeled M1-M10 for reference throughout the manuscript. The whole-pool diversity score (Step 1) was computed from M5-M7 and M10 (PSD, RSD, RStD, and PRF), whereas the benchmark composite score (Steps 2-3) was computed from all ten metrics (M1-M10). Here, $n$ denotes the benchmark size, $x_i$ the standardized feature vector of entry $i$, $d(x_i, x_j)$ the Euclidean distance between entries $i$ and $j$ in feature space, $S$ the benchmark sample, $P$ the corresponding non-redundant pool, $C(\cdot)$ the occupied feature-space bin, NND the nearest-neighbor distance, $s_i^{\text{pep}}$ and $s_i^{\text{rec}}$ the peptide and receptor sequences of entry $i$, respectively, $J_5$ the Jaccard similarity computed from receptor 5-mers, TM the pairwise receptor TM-score, $W_1$ the first Wasserstein (Earth Mover's) distance, and $K = \{\Delta G, \text{pep\_len}, \text{rec\_len}\}$ the set of feature dimensions used to evaluate distribution preservation.

| Label | ID | Metric | Definition |
|---|---|---|---|
| M1 | MNND | Mean nearest-neighbor distance | $\frac{1}{n}\sum_{i=1}^{n}\min_{j\neq i} d(x_i, x_j)$ |
| M2 | MPD | Mean pairwise distance | $\binom{n}{2}^{-1}\sum_{i<j} d(x_i, x_j)$ |
| M3 | FSC | Feature-space coverage | $\frac{\lvert C(S)\cap C(P)\rvert}{\lvert C(P)\rvert}$ |
| M4 | CVNND | Coefficient of variation of NND | $\frac{\sigma(\text{NND})}{\mu(\text{NND})}$ |
| M5 | PSD | Peptide sequence diversity | $1-\overline{\text{Identity}}$ |
| M6 | RSD | Receptor sequence diversity | $1-\overline{J_5}$ |
| M7 | RStD | Receptor structural diversity | $1-\overline{\text{TM}}$ |
| M8 | SCC | Structural cluster coverage | $\frac{\lvert Clusters(S)\rvert}{\lvert Clusters(P)\rvert}$ |
| M9 | WD | Normalized Wasserstein distance | $\frac{1}{\lvert K\rvert}\sum_{k\in K}\frac{W_1(P_k, S_k)}{\max(P_k)-\min(P_k)}$ |
| M10 | PRF | Peptide redundancy fraction | $\frac{\#\{(i,j):\text{Identity}>0.7\}}{\binom{n}{2}}$ |
